# Valorization of Paneer waste Whey through Fermentation with *Pediococcus pentosaceus* NCDC 273 Insights from Intracellular Metabolomics by GC-MS

**DOI:** 10.1101/2025.08.16.670691

**Authors:** Manish Singh Sansi, Daraksha Iram, Kamal Gandhi, Nishi Singh, Shiv Kumar Sood

**Affiliations:** Animal Biochemistry Division, ICAR-National Dairy Research Institute, Karnal, India; Dairy Microbiology Division, ICAR-National Dairy Research Institute, Karnal, India; Division of Dairy Chemistry, ICAR-National Dairy Research Institute, Karnal, India

**Author notes:** Corresponding author. Animal Biochemistry Division, ICAR-NDRI, India. Both are contributed equally to this work (Manish Singh Sansi, Daraksha Iram).

**Keywords:** GC-MS, dairy waste paneer whey, valorization, metabolites, *Pediococcus pentosaceus* NCDC 273, fermentation

## Abstract

The present study employed an untargeted GC-MS-based metabolomics approach to investigate the intracellular metabolic landscape of *Pediococcus pentosaceus* NCDC 273 during the fermentation of paneer whey. The goal was to understand the metabolic dynamics that enable this bacterium to efficiently utilize a dairy by-product and produce value-added compounds. Intracellular metabolites were profiled at the early (6 h) and late (18 h) exponential growth phases, revealing a distinct temporal metabolic reprogramming. Multivariate analysis and correlation networks confirmed significant metabolic shifts, where the early phase was characterized by active uptake and utilization of diverse sugars and fatty acids to support rapid proliferation. In contrast, the late phase demonstrated a metabolic reorientation towards energy production, redox balance, and stress adaptation. This was evidenced by the significant accumulation of key metabolites such as lactic acid, nicotinamide, and trehalose, which are crucial for maintaining growth and tolerance in an acidifying environment. Overall, the findings demonstrate that *P. pentosaceus* NCDC 273 possesses the metabolic flexibility to effectively channel whey-derived sugars into central metabolism, while concurrently deploying adaptive strategies for survival. This research represents the first intracellular metabolomic characterization of this strain during whey fermentation, providing novel mechanistic insights that reinforce its potential for the valorization of paneer whey into functional bioproducts.

## Introduction

The dairy industry is one of the largest contributors to global food production, generating an estimated 186 million tonnes (MT) of liquid whey annually, with an average growth rate of approximately 2% per year (USDEC, 2003). Despite its vast nutritional potential, whey is often considered an industrial by-product due to the lack of cost-effective treatment systems for its disposal. This has created a pressing challenge for the dairy sector, as the unregulated release of whey effluents into the environment not only contributes to water pollution but also represents a substantial loss of valuable nutrients. Whey typically contains about 55% of the original nutrients of milk, including 4.5% lactose, 0.8% soluble proteins, 1% mineral salts, and 0.1–0.8% lactic acid (Yang et al., 1994). Discarding such a nutrient-rich resource poses significant environmental and economic concerns, underlining the urgent need for sustainable and innovative utilization strategies.

Globally, researchers are exploring diverse biotechnological approaches to valorize whey. For instance, novel technologies such as ohmic heating and supercritical carbon dioxide treatment have been employed to develop whey-based beverages with improved physicochemical and sensory properties (Amaral et al., 2018; Cappato et al., 2018; Costa et al., 2018). Advances in fermentation technology have further enabled the conversion of whey lactose into high-value products like lactic acid using genetically engineered yeasts that can metabolize lactose efficiently (Turner et al., 2017). Lactic acid is particularly significant due to its wide range of industrial applications, including biodegradable plastics, pharmaceuticals, and food preservation, while also serving as a key metabolite in fermentation pathways (Roy et al., 1987). However, despite these advances, more than 50% of global whey production amounting to roughly 180 MT annually is still discarded, releasing nearly 1.5 MT of protein and 8.6 MT of lactose into the environment (Rai et al., 2020). This highlights the urgent necessity for sustainable solutions that integrate whey into value-added bioprocesses. Among microbial candidates for whey valorization, lactic acid bacteria (LAB) have garnered significant attention due to their metabolic versatility, probiotic properties, and ability to produce bioactive compounds. Lactiplantibacillus plantarum, a facultative heterofermentative LAB, is ubiquitous across diverse food matrices and the human microbiota. It is known for producing antimicrobial, antioxidant, and anti-inflammatory metabolites such as bacteriocins, cyclic dipeptides, and phenolic compounds (Bukhari et al., 2020; Mun et al., 2019; Salminen et al., 2021). Certain strains, such as L. plantarum HAC01 isolated from Korean kimchi, have demonstrated beneficial effects on host metabolism, fat accumulation, and gut microbiota modulation (Lee et al., 2021; Oh et al., 2021). Although several physiological benefits of LAB have been reported, the mechanistic understanding of their metabolic pathways and bioactive compound production remains incomplete. To address these knowledge gaps, metabolomics approaches particularly non-targeted and targeted metabolomics have emerged as powerful tools to comprehensively profile microbial metabolic outputs. Non-targeted metabolomics allows the unbiased detection of a wide array of metabolites, whereas targeted metabolomics provides sensitive and quantitative insights into specific compounds of interest (Gao et al., 2018; Teav et al., 2019). Mass spectrometry (MS)-based platforms, often coupled with chromatographic separation, have further enhanced the ability to capture diverse metabolites despite challenges associated with structural and chemical complexity (Tan et al., 2018; Tufi Lamoree et al., 2015). In this context, Pediococcus pentosaceus a homofermentative LAB species has emerged as a promising candidate for whey fermentation. P. pentosaceus not only exhibits efficient lactose-to-lactic acid conversion but also produces pediocin, a potent antimicrobial peptide with bio preservative potential (Oliveira et al., 2017; Garsa et al., 2014; Verma et al., 2019). Traditionally, P. pentosaceus has been applied in the fermentation of sausages, vegetables, and other foods due to its preservative properties (Papagianni & Anastasiadou, 2009; Smith & Palumbo, 1983). However, its potential for valorizing paneer whey into lactic acid and bioactive compounds remains largely unexplored. Therefore, the present study focuses on the utilization of paneer whey as a low-cost substrate for lactic acid production through fermentation by P. pentosaceus NCDC 273. By employing untargeted gas chromatography-mass spectrometry (GC–MS) profiling, we aim to investigate the metabolic changes associated with the growth of *P. pentosaceus* in paneer whey and assess its potential for lactic acid production. This work not only offers a sustainable solution to mitigate whey disposal issues but also highlights the dual benefits of generating high-value products such as lactic acid and bio-preservatives, thereby aligning with global efforts to promote circular bioeconomy in the dairy sector.

## 2. Material method

### 2.1. Chemicals and reagents

Paneer whey by-product samples were sourced from the National Dairy Research Institute, Karnal, and Haryana, India. Nutrient broth, agar, and other culture media components were procured from HiMedia, India. All chemicals used were of analytical grade. HPLC-grade solvents, including water, methanol, ethyl acetate, and acetonitrile, were obtained from Sigma Aldrich (Haryana, India). Formic acid, BSTFA [N,O-bis(trimethylsilyl)trifluoroacetamide], methoxyamine hydrochloride, pyridine, ammonium acetate, ammonium hydroxide, phosphate-buffered saline (PBS), and standard compounds were supplied by Sigma Aldrich (Haryana, India). De Man, Rogosa, and Sharpe (MRS) broth was also purchased from HiMedia, India.

### 2.2. Bacterial cultures and conditions

The lactic acid producing bacteria used for current study is *Pediococcus pentosaceus* (NCDC 273, National Collection of Dairy Cultures, National Dairy Research Institute, H.R., and India) available in our lab repository. The culture medium was used *Lactobacillus* deMan-Rogosa-Sharpe (MRS) broth (De Man et al., 1960). All isolates were stored in 50% glycerol stocks at -20°C.

### 2.3. Side stream waste whey fermented by *Pediococcus pentosaceus* NCDC 273

Paneer whey, a by-product of dairy processing, was utilized as a substrate for fermentation with *Pediococcus pentosaceus* NCDC 273 to generate and characterize bioactive metabolites through GC/MS analysis. Prior to large-scale fermentation, the strain was repeatedly sub-cultured and acclimatized in paneer whey to ensure stable growth. The whey medium was adjusted to pH 6.5 ± 0.2 at 25 °C using either 1 N HCl or 5 M NaOH and subsequently sterilized by autoclaving at 121.8 °C for 15 min. Fermentation was initiated by inoculating 1000 mL of sterilized whey contained in an Erlenmeyer flask with 25 mL of *P. pentosaceus* culture (ODLLL: 0.8–1.0), followed by incubation at 37 °C under shaking conditions (200 rpm) for 24 h. During the process, pH and cell density (ODLLL) were monitored as described by Liu et al. (2010). Aliquots of 100 mL were withdrawn at 12 and 24 h, after which the cultures were centrifuged at 11,000 × g for 10 min at 4 °C. The harvested cells were washed thrice with sterile 1X PBS and stored at −20 °C for subsequent intracellular metabolite extraction, following the protocol of Kim et al. (2023). At the completion of fermentation, the whey broth was first centrifuged at 4 °C and 1200 rpm for 10 min to remove residual debris. The supernatant was then subjected to a second centrifugation at 12,000 rpm for 10 min at 4 °C to collect the bacterial cells. The harvested cells from both groups were washed three times with sterile water and prepared for metabolomic profiling.

### 2.4. Non-targeted Intracellular analysis

#### 2.4.1. Sample extraction and preparation for GCMS analysis

Sample preparation of *Pediococcus pentosaceus* NCDC 273 was carried out with slight modifications from previously reported protocols (Yang et al., 2018; Li et al., 2012). The extraction solvent consisted of methanol, acetonitrile, and water mixed in the ratio 4:4:2 (v/v/v). Cell pellets obtained from 25 mL of culture were resuspended in 1 mL of pre-cooled (−20 °C) solvent, vortexed for 30 s, and subjected to three freeze–thaw cycles (liquid nitrogen for 5 min, followed by incubation at 4 °C for 20 min). The samples were then disrupted using sonication at a frequency of 30 sL¹ for 10 min (Meyer et al., 2010; Ming et al., 2018). After sonication, samples were centrifuged at 12,000 rpm for 10 min at 4 °C, and the resulting supernatants were transferred into clean tubes. Excess methanol was evaporated under nitrogen for about 2 h, after which the extracts were stored overnight at −40 °C. The samples were subsequently freeze-dried for ∼48 h and preserved in a desiccator until derivatization and analysis by GC–MS (Ming et al., 2018).

### 2.5. Derivatization of the bacterial intracellular metabolite for GCMS analysis

#### Derivatization procedure

For chemical derivatization, 200 μL of methoxyamine hydrochloride solution (15 mg/mL in pyridine) was added to each dried extract, vortexed for 30 s, and incubated at 37 °C for 90 min. Subsequently, 200 μL of N,O-bis(trimethylsilyl)trifluoroacetamide (BSTFA) containing 1% trimethylchlorosilane (TMCS) was introduced, followed by incubation at 70 °C for 30 min and an additional equilibration at room temperature for 30 min (Chen et al., 2014). After cooling, 100 μL of n-heptane containing phthalates (used as an internal standard) was added. The mixture was vortexed briefly and centrifuged at 12,000 rpm for 10 min at 4 °C. The resulting supernatant was combined with 800 μL of hexane, then transferred into silanized GC vials and stored until GC–MS analysis.

#### 2.5.1. GC-MS Analysis

Gas chromatography–mass spectrometry (GC–MS) was carried out using a Shimadzu GCMS-QP2010 Ultra system coupled with a GC-2010 unit and equipped with an AOC-20i+s autosampler (Shimadzu, Japan). A 1 µL aliquot of each derivatized sample was injected in split mode (split ratio 10:1) with helium as the carrier gas at a constant flow rate of 1.21 mL/min. The ion source temperature was maintained at 230 °C, while the interface was set to 270 °C. The solvent cut-off time was adjusted to 3.5 min, with acquisition in relative detector gain (ACQ) mode. Mass spectra were recorded over a scan range corresponding to 4.00–39.98 min, with an event time of 0.20 s and a scan speed of 3333. The oven temperature program began with an initial isothermal hold at 60 °C for 2 min, followed by a ramp of 10 °C/min up to 250 °C with a 5-min hold, and then further increased to 280 °C at 15 °C/min with a final hold of 12 min.

### 2.6. Data Processing and Statistical Analysis

Chromatographic peaks were processed using the GCMS Workstation LabSolution software (Shimadzu), and metabolite identification was achieved by comparison with the GCMS DB-Public-KovatsRI-VS3 mass spectral library with MSDIAL 5.52. The preliminary metabolite list was curated in Microsoft Excel to remove background contaminants. The refined dataset, including metabolite names and their peak areas, was exported as a CSV file and uploaded to the MetaboAnalyst 6.0 platform (https://www.metaboanalyst.ca/). Data pre-processing involved sum normalization followed by Pareto scaling. To discriminate among sample groups, supervised Partial Least Squares-Discriminant Analysis (PLS-DA) was applied. One-way ANOVA (p < 0.05). In addition, hierarchical clustering analysis (HCA) was conducted using Euclidean distance and Ward’s linkage to evaluate group-wise similarities. Only metabolites with an adjusted p-value < 0.05 were considered biologically significant. All experiments were carried out with four biological replicates (n = 4), and results were presented as mean ± standard deviation.

## 3. Results and Discussion

The present study investigated the intracellular metabolic landscape of *Pediococcus pentosaceus* NCDC 273 during the fermentation of paneer whey, a dairy industry by-product traditionally regarded as waste but increasingly recognized as a valuable substrate for microbial biotransformation. Using an untargeted GC–MS-based metabolomics approach, the intracellular metabolites of *P. pentosaceus* were profiled at different growth stages in supplemented paneer whey medium (SPWM). Consistent with our earlier findings and previous reports, *P. pentosaceus* NCDC 273 displayed rapid exponential growth in SPWM, reaching approximately 9.1 log cfu mlL¹ within 24 h, accompanied by a decline in pH from 6.97 to 4.25 due to lactic acid accumulation (Verma et al., 2023; Fu & Mathews, 1999). HPLC analysis confirmed a fermentation pattern, with optimal proliferation observed at moderately acidic conditions (pH 4.98–6.97), while growth was strongly inhibited under highly acidic environments (pH ∼3.98). The fermentation process was nearly complete within 36 h, which agrees with earlier reports of maximum growth occurring between 20 and 24 h in glucose- and lactose-based media (Simha et al., 2012; Verma et al., 2023; Liu et al., 2025).

To capture the metabolic shifts during fermentation, cell pellets collected at 6 h (early exponential phase) and 18 h (late exponential phase) were analyzed for intracellular metabolites. The GC–MS analysis enabled the detection of 59 distinct metabolites, spanning central metabolic pathways showed in (Table 1). These metabolites were predominantly associated with carbohydrate metabolism, amino acid biosynthesis and degradation, nucleotide metabolism, and lipid-related pathways. Overall, the metabolomic data provide evidence that *P. pentosaceus* NCDC 273 effectively channels whey-derived sugars into central metabolism and lactic acid production while maintaining metabolic flexibility under stress conditions. The accumulation of stress-related metabolites reflects the strain’s capacity to tolerate acidification, a key feature for its survival and efficiency in dairy fermentation environments. Importantly, these findings represent the first intracellular metabolomic characterization of *P. pentosaceus* NCDC 273 during whey fermentation, offering novel insights into its metabolic potential and reinforcing its applicability for the valorization of paneer whey into functional, value-added bioproducts.

**Table 1.** List of intracellular metabolites identified based on untargeted metabolomics analysis.

| Query | Match | HMDB |
| --- | --- | --- |
| Hexacosanoic acid | Hexacosanoic acid | HMDB0002356 |
| Xylonic acid | D-Xylonic acid | HMDB0059750 |
| Valine | L-Valine | HMDB0000883 |
| Uridine | Uridine | HMDB0000296 |
| Urea | Urea | HMDB0000294 |
| Uridine 5'-monophosphate | Uridine 5'-monophosphate | HMDB0000288 |
| Trehalose | Trehalose | HMDB0000975 |
| Threonine | L-Threonine | HMDB0000167 |
| Tetracosane | Lignocerane | HMDB0034282 |
| Tartaric acid | Tartaric acid | HMDB0000956 |
| Sucrose-6-phosphate | Sucrose-6-phosphate | HMDB0258553 |
| Sucrose | Sucrose | HMDB0000258 |
| Succinic acid | Succinic acid | HMDB0000254 |
| Stearic acid | Stearic acid | HMDB0000827 |
| Shikimic acid | Shikimic acid | HMDB0003070 |
| Serine | Serine | HMDB0000187 |
| Pyridoxamine 5'-phosphate | Pyridoxamine 5'-phosphate | HMDB0001555 |
| Pseudouridine | Pseudouridine | HMDB0000767 |
| Proline | Proline | HMDB0000162 |
| Phosphate | Phosphate | HMDB0001429 |
| Palmitic acid | Palmitic acid | HMDB0000220 |
| Nicotinamide | Niacinamide | HMDB0001406 |
| Myristic acid | Myristic acid | HMDB0000806 |
| Myo-Inositol | myo-Inositol | HMDB0000211 |
| Methylmalonic acid | Methylmalonic acid | HMDB0000202 |
| Melibiose | Melibiose | HMDB0000048 |
| Melezitose | Melezitose | HMDB0011730 |
| Maltotriose | Maltotriose | HMDB0001262 |
| Maltose | D-Maltose | HMDB0000163 |
| Maltitol | Maltitol | HMDB0002928 |
| L-Serine | Serine | HMDB0000187 |
| Linolenic acid | alpha-Linolenic acid | HMDB0001388 |
| Linoleic acid | Linoleic acid | HMDB0000673 |
| L-Iditol | L-Iditol | HMDB0011632 |
| Leucine | Leucine | HMDB0000687 |
| Lactulose | Lactulose | HMDB0000740 |
| Lactose | Alpha-Lactose | HMDB0000186 |
| Lactitol | Lactitol | HMDB0040937 |
| Lactic acid | Lactic acid | HMDB0000190 |
| Inositol | myo-Inositol | HMDB0000211 |
| Icosanoic acid | Arachidic acid | HMDB0002212 |
| Heptadecanoic acid | Heptadecanoic acid | HMDB0002259 |
| Glycerol | Glycerol | HMDB0000131 |
| Galactinol | Galactinol | HMDB0005826 |
| Fructose-2,6-bisphosphate | D-Fructose 2,6-bisphosphate | HMDB0001047 |
| D-Ribose 5-phosphate | D-Ribose 5-phosphate | HMDB0001548 |
| D-Raffinose | Raffinose | HMDB0003213 |
| D-Pantothenic acid | Pantothenic acid | HMDB0000210 |
| Docosenoic acid | Erucic acid | HMDB0002068 |
| D-Melezitose | Melezitose | HMDB0011730 |
| D-Mannitol | Mannitol | HMDB0000765 |
| D-Galactosamine | D-Galactosamine | METPA0263 |
| D-Fructose 6-phosphate | Fructose 6-phosphate | HMDB0000124 |
| D-Fructose | D-Fructose | HMDB0000660 |
| Citric acid | Citric acid | HMDB0000094 |
| Cerotinic acid | Hexacosanoic acid | HMDB0002356 |
| Behenic acid | Behenic acid | HMDB0000944 |
| 2-Deoxyribose 5-phosphate | Deoxyribose 5-phosphate | HMDB0001031 |
| 1-Kestose | 1-Kestose | HMDB0011729 |

### 3.3 Metabolite distribution before and after normalization

The intracellular metabolite profiling of *Pediococcus pentosaceus* NCDC 273 grown in paneer whey at 6 h (S1–S1.3) and 18 h (S2–S2.3) revealed distinct metabolic shifts (Figure 1). Before normalization, metabolite concentrations exhibited high variability, with only a few metabolites, such as lactose, trehalose, uridine, and phosphate, showing elevated levels. After normalization, the distribution was balanced, allowing clearer visualization of metabolite dynamics. At 6 h, fatty acids (linoleic acid, palmitic acid), disaccharides (lactose, trehalose), citric acid, and uridine were relatively abundant, indicating early substrate utilization and initial energy metabolism. In contrast, at 18 h, significantly higher levels of nicotinamide, phosphate, glycolytic intermediates (D-fructose-6-phosphate, shikimic acid), and lactic acid were observed, reflecting enhanced bacterial growth, active glycolysis, and lactic acid fermentation. These results demonstrate a time-dependent intracellular metabolic reprogramming, where *P. pentosaceus* shifts from substrate utilization during early growth (6 h) to energy production and redox balance during advanced fermentation (18 h) shown in Figure 1.

**Figure 1.**
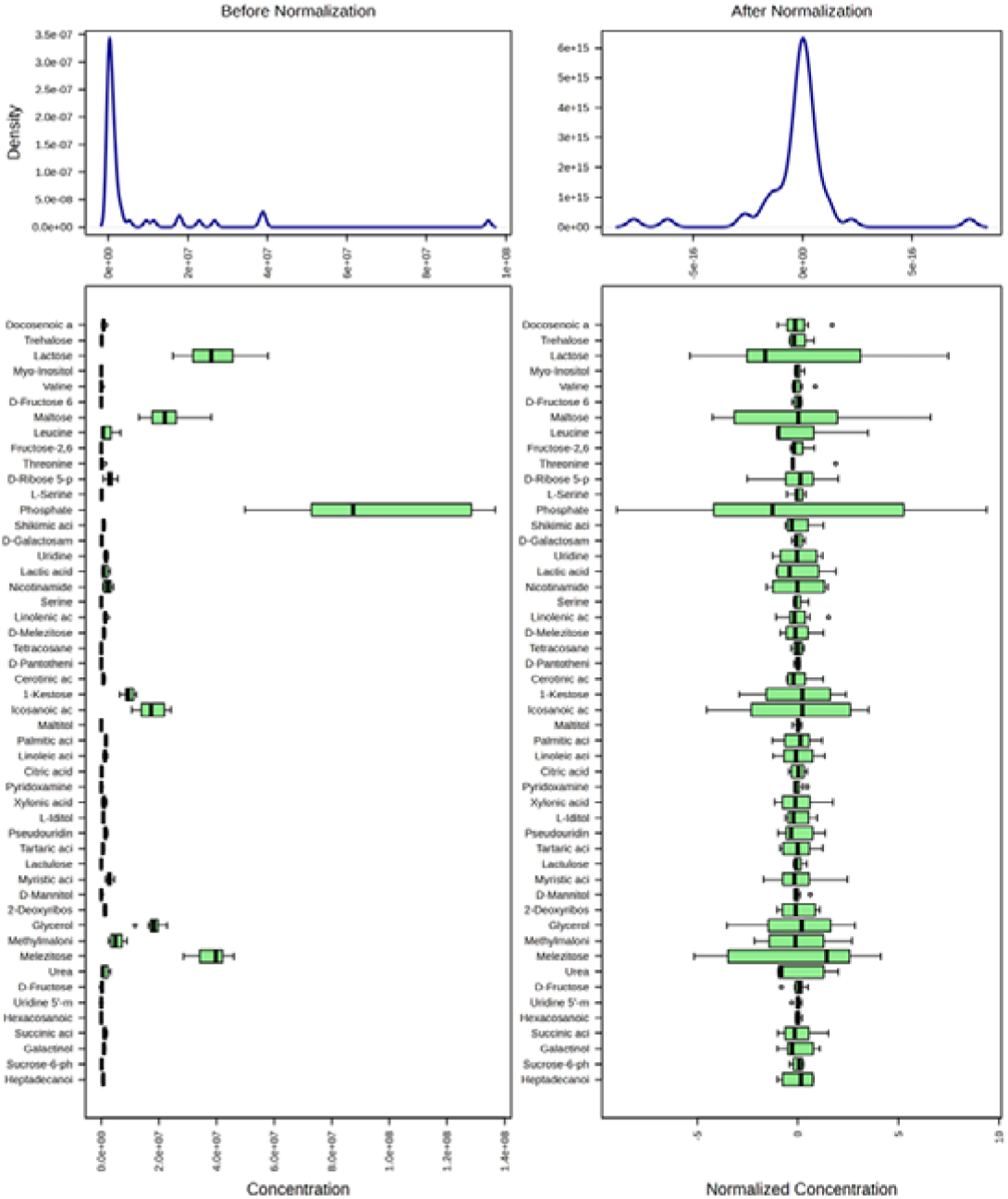
Intracellular metabolites of *Pediococcus pentosaceus* NCDC 273 grown in paneer whey at 6 h (S1–S1.3) and 18 h (S2–S2.3). Density plots (top) and box plots (bottom) show metabolite distribution before and after normalization. Normalization reduced variability and enabled clearer comparison of metabolite profiles across time points.

### 3.4. Volcano plot showing differential intracellular metabolites of *P. pentosaceus* NCDC 273

The intracellular metabolite profile of *Pediococcus pentosaceus* NCDC 273 grown in paneer whey revealed distinct differences between 6 h and 18 h of incubation. The volcano plot demonstrated that several metabolites were significantly altered over time, highlighting dynamic metabolic adaptation during bacterial growth. At 18 h, metabolites such as nicotinamide and uridine were markedly upregulated, indicating enhanced activity of nucleotide metabolism and NADL/NADH-related redox balance, which are essential for sustaining bacterial growth and energy metabolism in later stages of fermentation. Similarly, the accumulation of heptadecanoic acid suggests alterations in fatty acid metabolism, possibly linked to membrane remodeling and energy storage during prolonged growth. In contrast, pyridoxamine 5′-phosphate and proline were significantly downregulated at 6 h, suggesting their early utilization in amino acid metabolism and coenzyme-dependent reactions to support initial bacterial proliferation showed in (Figure 2). Collectively, these findings suggest that *P. pentosaceus* undergoes a time-dependent metabolic reprogramming in paneer whey, with early consumption of amino acids and cofactors followed by enhanced synthesis of nucleotides, fatty acids, and redox-related metabolites to sustain growth and metabolic activity at later stages. Zhang et a., (2025) revealed study that, the result of volcano plot showed significant metabolic reprogramming in LPFJ after fermentation, with 28 metabolites up-regulated and 23 down-regulated (*p* < 0.05, FC ≥ 1.2). Up-regulated compounds included pyridoxine, indole acetaldehyde, 5-L-glutamyl-L-alanine, and 5-hydroxy-indole-acetaldehyde, indicating enhanced amino acid metabolism and increased vitamin B6 biosynthesis. In contrast, xanthosine, vanilloside, and D-glucarate were down-regulated, reflecting reduced activity in nucleotide and carbohydrate metabolism. These findings suggest that fermentation enriches bioactive metabolites while down-regulating certain energy and nucleotide intermediates, thereby improving the nutritional and functional potential of LPFJ.

**Figure 2.**
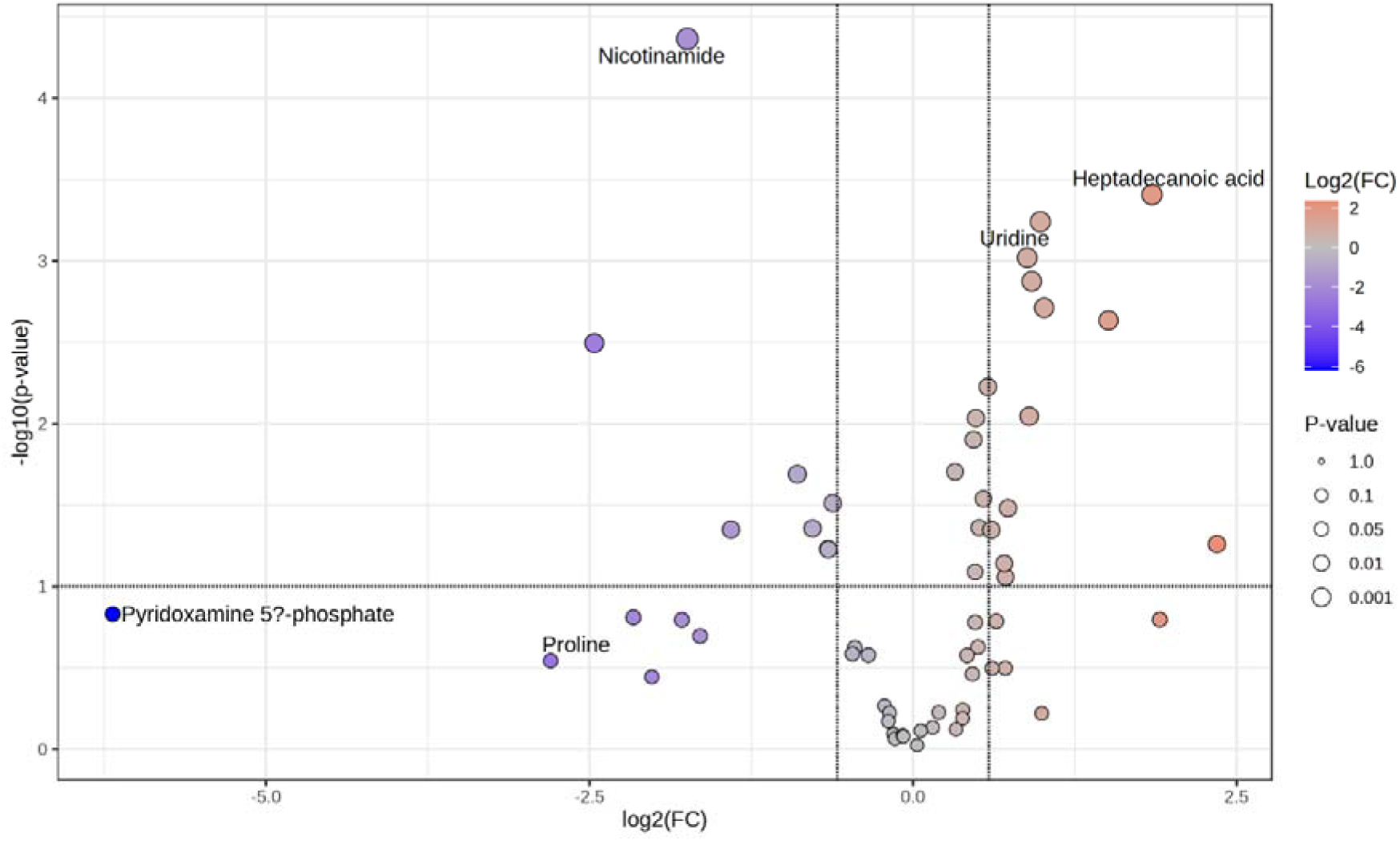
Volcano plot showing differential intracellular metabolites of *Pediococcus pentosaceus* NCDC 273 grown in paneer whey at 6 **h and 18 h**. The x-axis represents log2 fold change (FC), while the y-axis indicates –log10 (p-value). Red-colored metabolites indicate significant upregulation, whereas blue-colored metabolites indicate downregulation. Key metabolites such as nicotinamide, uridine, and heptadecanoic acid were upregulated at 18 h, while pyridoxamine 5′-phosphate and proline were downregulated at 6 h.

### 3.5. Intracellular metabolomic variability

The PCA score plot of intracellular metabolites obtained from *Pediococcus pentosaceus* NCDC 273 during SPWM fermentation at 6 h and 18 h showed a clear separation of the two groups, indicating distinct temporal metabolic reprogramming (Figure 3a,b). The first principal component (PC1) explained 60.2% of the total variance, while PC2 accounted for 18.6%, together capturing 78.8% of the metabolic variability. Samples corresponding to 6 h (pink cluster) were tightly grouped and distinctly separated from those of 18 h (green cluster), highlighting both reproducibility and significant differences in intracellular metabolic states at the two time points shown in (Figure 3a).

**Figure 3.**
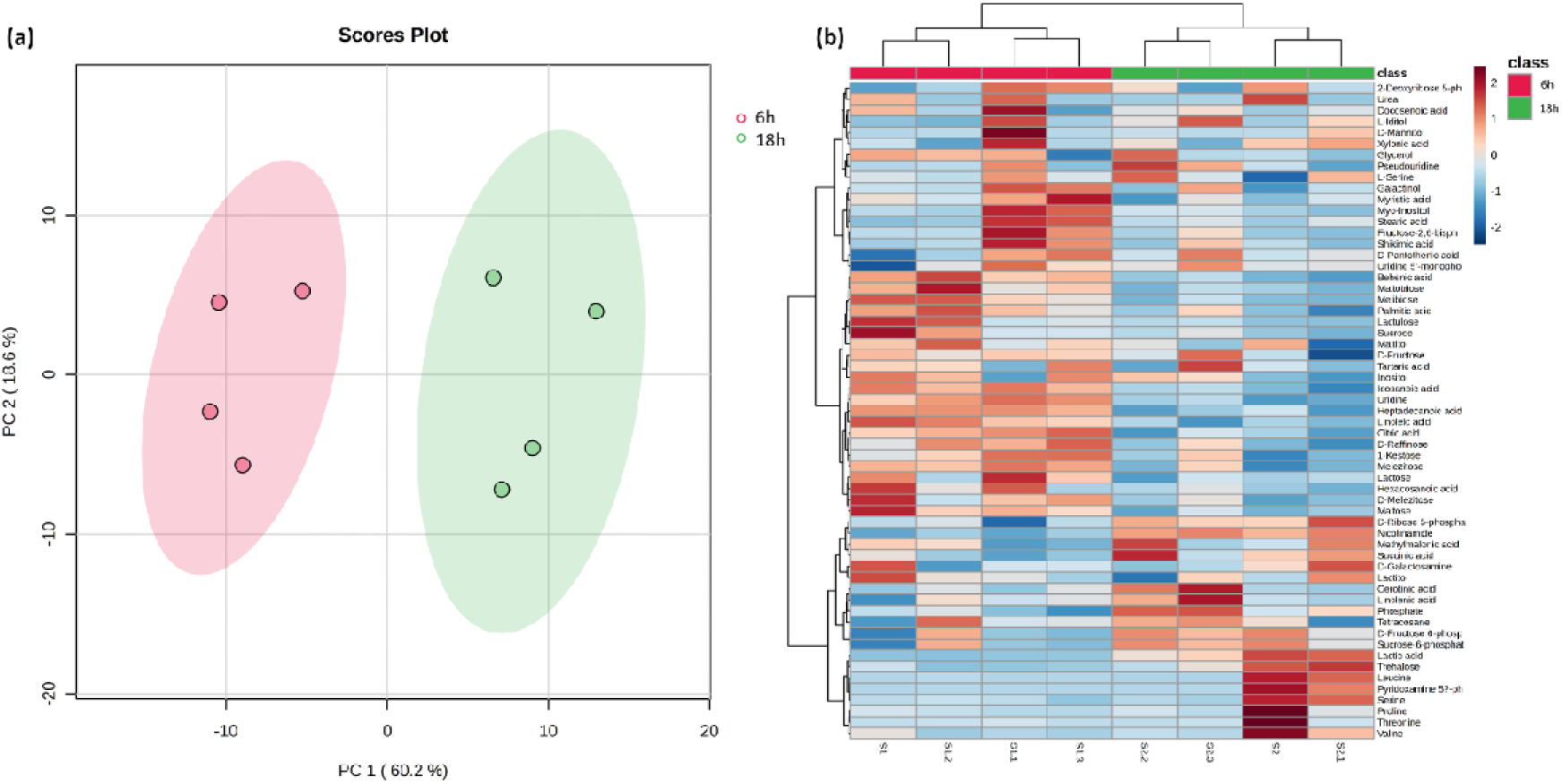
Multivariate analysis of intracellular metabolites during paneer whey fermentation by *Pediococcus pentosaceus* NCDC 273 at 6 h and 18 h. (A) Principal Component Analysis (PCA) score plot showing clear separation between 6 h and 18 h samples, indicating distinct metabolic shifts during fermentation. Replicates within each group clustered tightly, demonstrating good reproducibility, while PC1 and PC2 captured the major sources of variance related to fermentation time. (B) Hierarchical clustering heat map of significantly altered metabolites, where red and blue represent higher and lower relative abundances, respectively. Distinct metabolite groups were differentially regulated between 6 h and 18 h, highlighting dynamic intracellular metabolic reprogramming during the fermentation process.

The heat map analysis revealed distinct changes in the intracellular metabolite profiles of SPWM fermented by *Pediococcus pentosaceus* NCDC 273 at 6 h and 18 h. At 6 h, several metabolites such as 2-deoxyribose-5-phosphate, urea, D-mannitol, pseudouridine, L-serine, galactinol, and fatty acids (myristic acid, stearic acid, palmitic acid) were significantly upregulated, indicating active carbohydrate and amino acid metabolism during the early phase of fermentation. In contrast, the 18 h samples showed a marked accumulation of organic acids (citric acid, succinic acid, lactic acid), short-chain fatty acids (linoleic acid, hexacosanoic acid), and sugar derivatives (raffinose, maltose, sucrose, fructose-6-phosphate, sucrose-6-phosphate), suggesting a metabolic shift towards enhanced energy production and acidogenesis at the later fermentation stage. Cluster analysis further demonstrated a clear separation between 6 h and 18 h groups, confirming that the intracellular metabolome of *P. pentosaceus* underwent significant temporal reprogramming. The early stage (6 h) was dominated by intermediates of nucleotide and amino acid metabolism, while the late stage (18 h) reflected increased glycolytic flux, TCA cycle intermediates, and biosynthesis of fatty acids and organic acids shown in (Figure 3b).

### 3.6. Intracellular metabolic profile difference between 6 and 18 h

The box plot analysis of intracellular metabolites during 6 h and 18 h fermentation of paneer whey by *P. pentosaceus* NCDC 273 revealed significant temporal metabolic reprogramming. Lactic acid bacteria (LAB) require sugars as their primary energy source for growth and metabolism (Bintsis et al., 2018). To facilitate carbohydrate uptake, they employ multiple transport systems, including phosphotransferase systems (PTS), ATP-binding cassette (ABC) transporters, permeases, and symporters (Thompson et al., 1988). In the present study, a high intracellular accumulation of lactose was observed in *P. pentosaceus* NCDC 273, likely due to its rapid lactose uptake from the medium (Figure 4). This extensive lactose uptake may contribute to the rapid growth rates characteristic of NCDC 273 strains (Verma et al., 2023). At 6 h of fermentation, higher intracellular levels of linoleic acid, maltose, myristic acid, sucrose, uridine, behenic acid, citric acid, heptadecanoic acid, D-melezitose, icosanoic acid, lactose, and lactulose were detected. These metabolites largely represent fatty acids, energy storage compounds, and nucleotide precursors, suggesting active uptake and biosynthetic processing of paneer whey constituents during the early exponential growth phase. Such accumulation reflects the cellular demand for macromolecular building blocks to sustain rapid biomass production. Consistent with this, homofermentative LAB such as *Lactococcus* and *Lactiplantibacillus* species utilize carbohydrates primarily via the Embden–Meyerhof– Parnas pathway, yielding lactic acid and ATP to support energy-intensive growth (McDonald et al., 1987; Wang et al., 2021).

**Figure 4.**
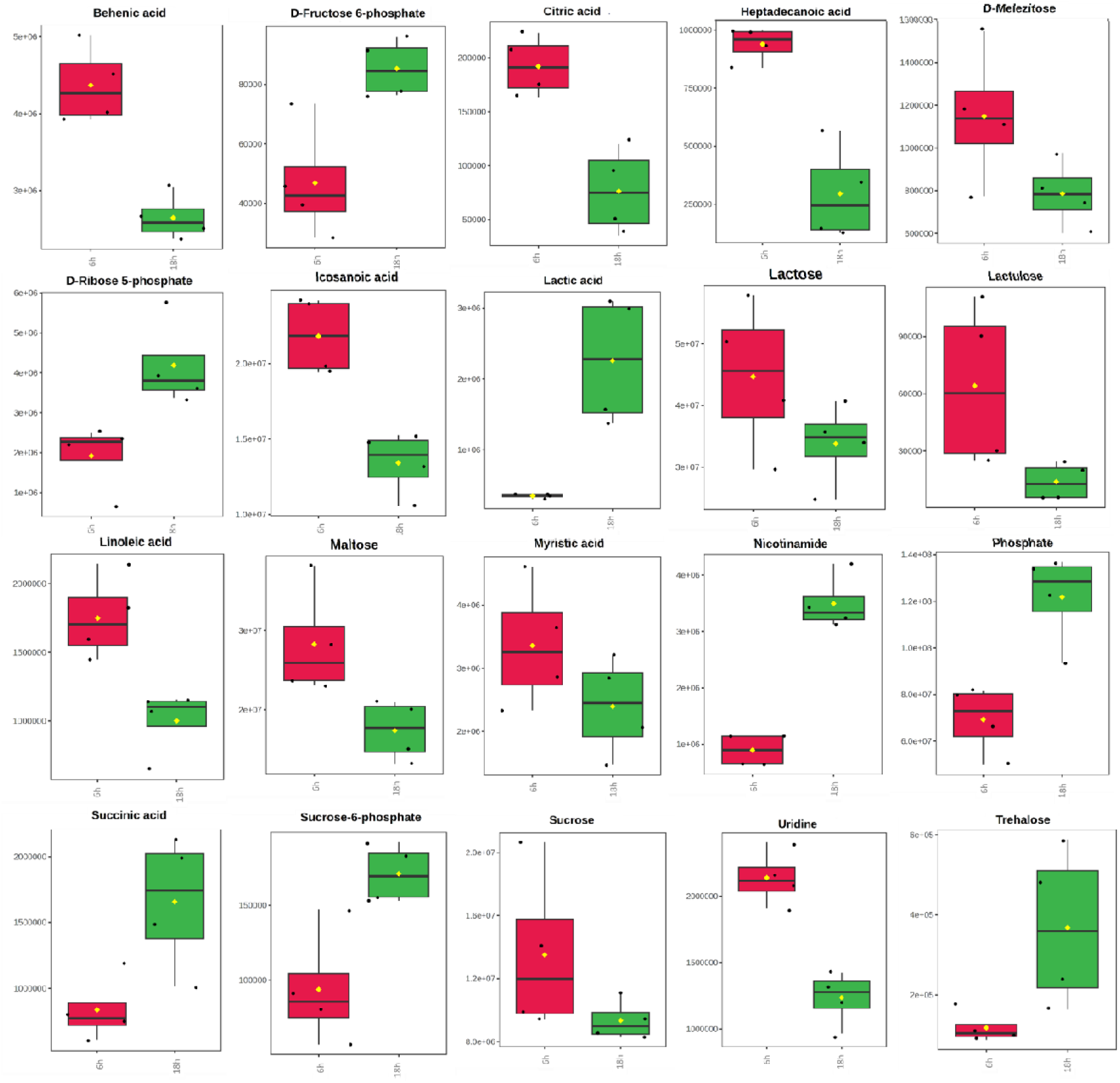
Box plots showing intracellular metabolite distribution during paneer whey fermentation by *Pediococcus pentosaceus* NCDC 273. At 6 h, higher levels of fatty acids, disaccharides, citric acid, and uridine were detected, indicating early substrate utilization. At 18 h, increased levels of nicotinamide, phosphate, glycolytic intermediates, and lactic acid were observed, reflecting enhanced metabolic activity and fermentation progression.

In contrast, the metabolite profile at 18 h indicated a substantial shift in intracellular composition. Elevated levels of nicotinamide, phosphate, succinic acid, sucrose-6-phosphate, trehalose, D-fructose-6-phosphate, D-ribose-5-phosphate, and lactic acid were observed. These metabolites are associated with glycolysis, the pentose phosphate pathway, nucleotide biosynthesis, and fermentation end products, highlighting a metabolic reorientation toward energy generation, redox balance, and stress adaptation during prolonged fermentation. In particular, the increased accumulation of lactic acid and succinic acid confirmed sustained carbohydrate fermentation, while elevated nicotinamide and phosphate intermediates indicated enhanced requirements for cofactors and energy metabolism at this stage. Simultaneously, the marked depletion of maltose, sucrose, and lactose between 6 h and 18 h underscored efficient sugar utilization by *P. pentosaceus*. Interestingly, significant intracellular accumulation of trehalose and maltose was detected during the mid-stationary phase, aligning with previous findings in *Lactobacillus sakei* (Fu et al., 2022; Lee et al., 2017). Beyond serving as a carbon source, trehalose is well documented for its role in protecting LAB against environmental fluctuations such as freeze–thaw or heat–cooling cycles (Mizunoe et al., 2018; Vaessen et al., 2019). Thus, trehalose accumulation by *P. pentosaceus* NCDC 273 may contribute not only to metabolic flexibility but also to stress resilience during extended fermentation.

Overall, the box plot data clearly demonstrate a dynamic metabolic transition between early (6 h) and late (18 h) fermentation stages. The early phase is dominated by substrate uptake and biosynthesis, while the late phase reflects central carbon metabolism activation, redox homeostasis, and adaptive stress responses. These results highlight the metabolic versatility of *P. pentosaceus* NCDC 23 in utilizing paneer whey as a growth substrate and provide mechanistic insights into its potential application in functional dairy fermentations.

### 3.7. Metabolic changes over time and VIP score

The biplot analysis revealed a clear separation between the metabolic profiles of *Pediococcus pentosaceus* NCDC 273 at 6 h and 18 h of growth in SPWM, indicating a distinct temporal shift in the intracellular metabolome. The red cluster (6 h) and green cluster (18 h) were clearly segregated along Principal Component 1 (PC1), which accounted for 60.2% of the total variance, highlighting it as the dominant factor driving metabolic differentiation (Figure 5a). Such separation underscores significant reprogramming of the metabolic state as the culture progressed from early exponential to mid-stationary phase. Among the metabolites, phosphate emerged as the most critical biomarker for distinguishing the two time points, with a VIP score approaching 4.0. Its strong association with the 18 h samples suggested a substantial increase in intracellular phosphate levels as the culture matured. Phosphate plays a central role in bacterial metabolism, particularly in *Lactococcus* species such as *Lactococcus lactis*, where it is vital for glycolysis, growth rate, and cellular biomass production (Antwi et al., 2008). The elevated phosphate levels in our study indicate a metabolic shift toward processes linked to energy regulation and structural maintenance during the stationary phase. In contrast, several sugars including D-Lactose, D-Maltose, Melibiose, Melezitose, and Maltotriose were dominant at the 6 h time point, reflecting their active utilization during exponential growth. This observation is consistent with the metabolic behavior of homofermentative LAB such as *Lactococcus* and *Lactiplantibacillus* spp., which preferentially metabolize carbohydrates through the Embden–Meyerhof–Parnas (EMP) pathway, yielding lactic acid and ATP to sustain rapid biomass accumulation (McDonald et al., 1987; Wang et al., 2021). Interestingly, trehalose and maltose accumulated significantly during the mid-stationary phase, aligning with previous reports in *Lactobacillus sakei*, where intracellular sugar accumulation was linked to stress adaptation and stationary phase physiology (Lee et al., 2017; Fu et al., 2022).

**Figure 5.**
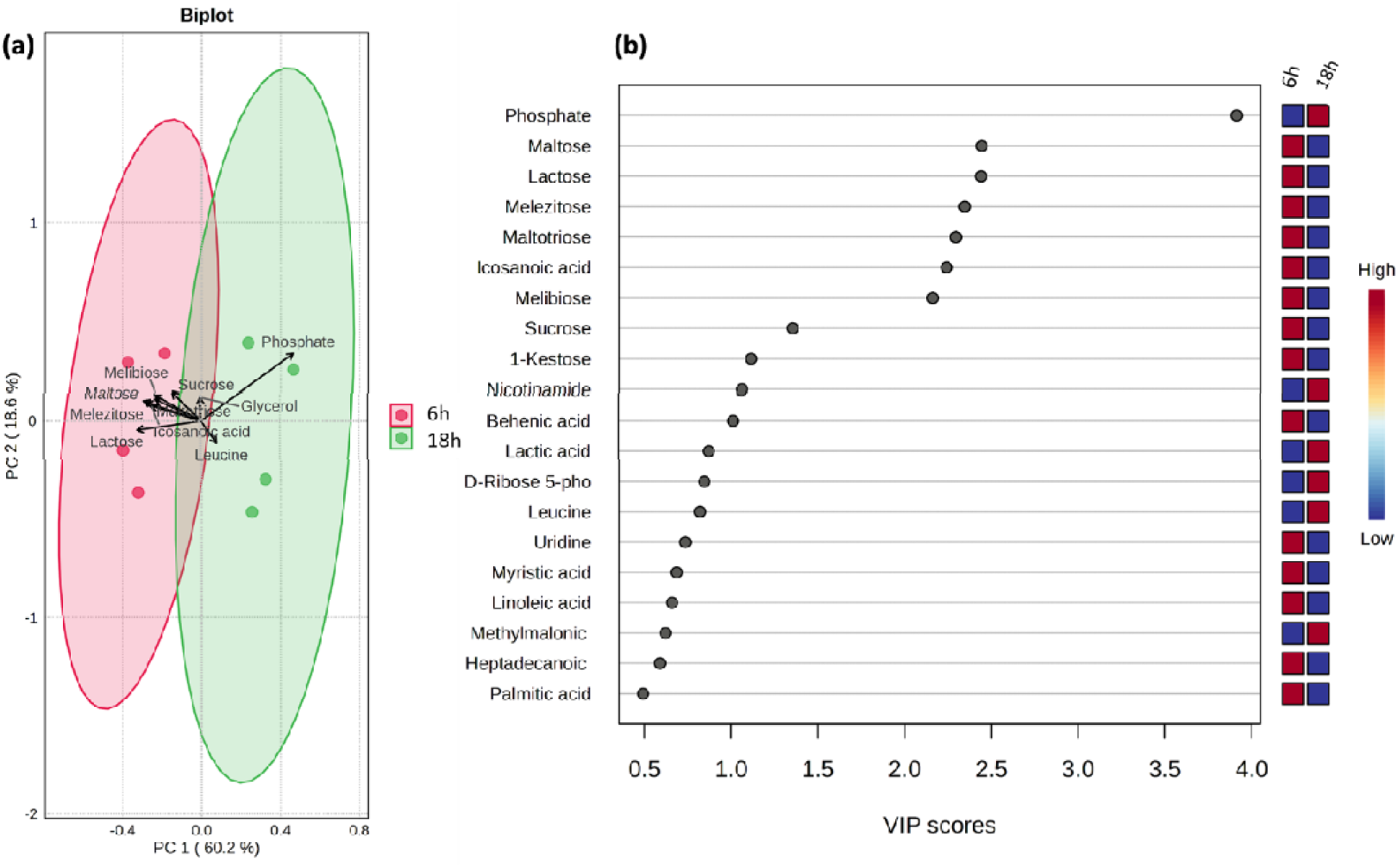
(a) Principal Component Analysis (PCA) biplot of the intracellular metabolic profile. (b) The primary drivers of this change are a few key metabolites, as highlighted by their high **Variable Importance in Projection (VIP)** scores.

The VIP score plot further reinforced these findings, with D-Maltose and D-Lactose ranked as the second and third most influential metabolites (VIP ≈ 2.5), confirming their role in distinguishing growth phases. Melibiose, Melezitose, and Maltotriose also showed significant contributions, indicating dynamic changes in their intracellular levels between 6 h and 18 h (Figure 5b). Together, these results suggest that the metabolic transition from 6 h to 18 h in *P. pentosaceus* NCDC 273 involves a shift from rapid sugar consumption to a phase characterized by phosphate accumulation and increased levels of metabolites such as glycerol and icosanoic acid. This shift reflects an adaptation strategy where the bacteria transition from energy-intensive growth toward alternative metabolic pathways that likely support stress tolerance, energy storage, and long-term survival in the paneer whey environment.

### 3.8. Network analysis of intracellular metabolites

The metabolite–metabolite interaction network analysis provided further insights into the intracellular metabolic organization of *P. pentosaceus* NCDC 273 during growth in paneer whey. The correlation network (Figure 6) revealed distinct clusters of positively and negatively correlated metabolites, suggesting coordinated metabolic processes as well as antagonistic relationships between specific metabolites. A prominent feature of the network was the strong positive correlations (red lines) observed among sugars and their related metabolites. For instance, sucrose was positively correlated with lactulose and lactose, both of which also showed positive correlations with melibiose and maltotriose. This pattern suggests that these sugars are tightly linked in the metabolic pathways of *P. pentosaceus* and are likely consumed or produced in a coordinated manner during growth. Such clustering of sugars is consistent with their role as primary energy sources in lactic acid bacteria, where multiple carbohydrates can be metabolized simultaneously through overlapping pathways.

**Figure 6.**
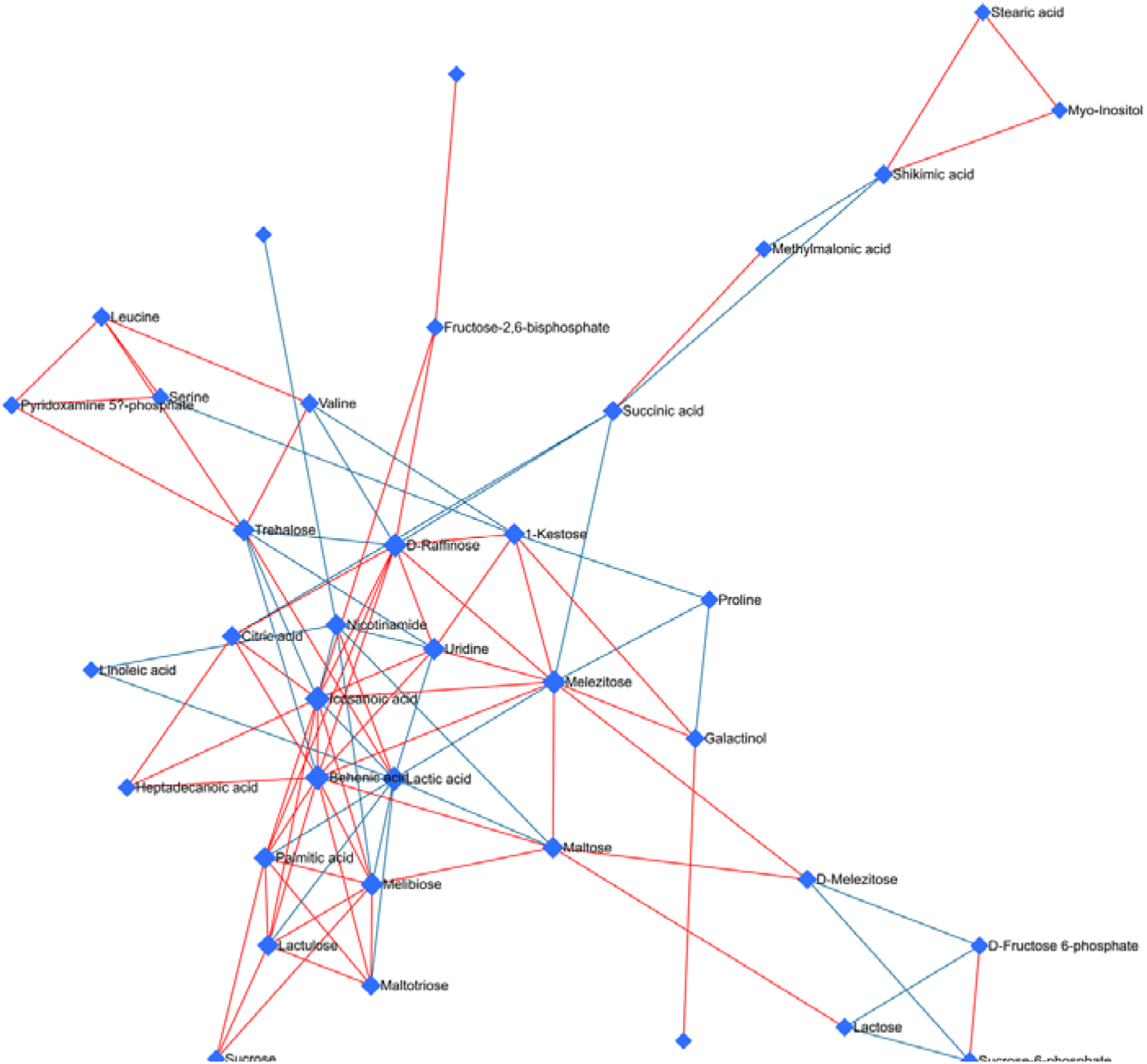
Network of intracellular metabolites from *Pediococcus pentosaceus* NCDC 273. The nodes (blue diamonds) represent individual metabolites. The lines between the nodes represent correlations in their concentrations. Red lines indicate a positive correlation, where the concentrations of both metabolites tend to increase or decrease together. Blue lines indicate a negative correlation, showing an inverse relationship. This network highlights the complex relationships and metabolic pathways within the bacteria, such as the positive correlations among various sugars and fatty acids, and the negative correlations between different metabolic clusters.

Conversely, the network revealed several negative correlations (blue lines), indicating antagonistic relationships between metabolites. For example, fructose-2,6-bisphosphate was negatively correlated with succinic acid, suggesting differential routing of carbon flux depending on growth stage or metabolic demand. Similarly, amino acid metabolism showed contrasting correlations: while leucine and serine were positively correlated, leucine displayed a negative correlation with valine. This suggests that although leucine and serine may share cooperative metabolic regulation, leucine and valine likely follow divergent metabolic fates, possibly reflecting differences in branched-chain amino acid metabolism and regulation.

Such correlation-based metabolite networks provide a systems-level view of metabolic coordination and competition within microbial cells. Similar approaches have been employed in plant systems, where metabolite–metabolite interaction networks were shown to highlight functional relationships between annotated metabolites in *Thompson Seedless* grapes (Jadhav et al., 2021). By analogy, the present findings underscore the metabolic flexibility of *P. pentosaceus* and highlight how sugars, organic acids, and amino acids are tightly regulated during different phases of growth. Overall, the network analysis emphasizes that while sugars such as sucrose, lactose, and melibiose form a cooperative metabolic cluster, other metabolites such as fructose-2,6-bisphosphate and succinic acid exhibit antagonistic relationships, reflecting dynamic shifts in central metabolism. These findings contribute to a deeper understanding of metabolic coordination in *P. pentosaceus* and suggest potential regulatory nodes that may control the balance between carbohydrate utilization and amino acid metabolism during growth in whey.

## Conclusion

This study provides the first intracellular metabolomic characterization of *Pediococcus pentosaceus* NCDC 273 during paneer whey fermentation using an untargeted GC-MS approach. The temporal profiling revealed a clear metabolic transition from active sugar and fatty acid utilization in the early phase to enhanced energy generation, redox balance, and stress adaptation in the late phase. The pronounced accumulation of lactic acid confirmed efficient fermentation of whey-derived lactose, while elevated levels of nicotinamide and trehalose highlighted metabolic strategies that support energy requirements and survival in an acidifying environment. These findings underscore the metabolic flexibility of *P. pentosaceus* NCDC 273 and its ability to reprogram intracellular metabolism in response to substrate availability and stress. Overall, the study demonstrates the strain’s potential for the sustainable valorization of paneer whey into value-added functional bioproducts, thereby contributing to both waste management and biotechnological innovation.

## Notes

### Competing Interest Statement

The authors have declared no competing interest.

